# BirdVox: Machine listening for bird migration monitoring

**DOI:** 10.1101/2022.05.31.494155

**Authors:** Vincent Lostanlen, Aurora Cramer, Justin Salamon, Andrew Farnsworth, Benjamin M. Van Doren, Steve Kelling, Juan Pablo Bello

## Abstract

The steady decline of avian populations worldwide urgently calls for a cyber-physical system to monitor bird migration at the continental scale. Compared to other sources of information (radar and crowdsourced observations), bioacoustic sensor networks combine low latency with a high taxonomic specificity. However, the scarcity of flight calls in bioacoustic monitoring scenes (below 0.1% of total recording time) requires the automation of audio content analysis. In this article, we address the problem of scaling up the detection and classification of flight calls to a full-season dataset: 6672 hours across nine sensors, yielding around 480 million neural network predictions. Our proposed pipeline, BirdVox, combines multiple machine learning modules to produce per-species flight call counts. We evaluate BirdVox on an annotated subset of the full season (296 hours) and discuss the main sources of estimation error which are inherent to a real-world deployment: mechanical sensor failures, sensitivity to background noise, misdetection, and taxonomic confusion. After developing dedicated solutions to mitigate these sources of error, we demonstrate the usability of BirdVox by reporting a species-specific temporal estimate of flight call activity for the Swainson’s Thrush *(Catharus ustulatus)*.

## Introduction

### The role of bioacoustic monitoring in conservation science

The northeast United States has lost 13% of its overall avian migratory biomass between 2007 and 2017 [1]. In this respect, the order *(Passeriformes)* appears as most direly affected: the global populations of American sparrows *(Passerellidae)* and New World warblers *(Parulidae)* have both declined by about 30% since 1970; i.e., 1.5 billion missing individuals. A large portion of this decline is attributable to human activity. The three biggest causes to avian mortality in the United States are predation by domestic cats, collisions with buildings, and collisions with vehicles [2]. Beyond these direct threats, global warming accelerates the risk of extinction in the near future: over two thirds of North American birds will become moderately or highly vulnerable to climate change under a 3.0 °C scenario [3].

American sparrows and New World warblers are both families that include long-distance migrants, with typical endpoints being Canada and Central America [4]. During migration, birds may alter their routes depending on local weather and availability of foraging resources [5]. For this reason, the number of individuals flying over any given area may vary from one year to the next and cannot be extrapolated from past observations alone. Instead, avian conservation science requires tools to monitor bird migration with a low latency of a few minutes at best to a few days at most [6]. The prospect of forecasting the quantity of vulnerable species of birds near hazardous sites (e.g., airports [7], windfarms [8], and dense urban areas [9]) creates an opportunity for preventive measures at the local level, such as the temporary reduction of light pollution [10]. Furthermore, the design of a low-latency and reliable system for bird migration monitoring would benefit civil aviation safety and agricultural planning [11].

At present, the two most widespread methods to monitor bird migration over time are direct human observations and remote sensing by radar [12]. Although these methods are scalable and informative, they exemplify a tradeoff between selection bias and taxonomic uncertainty. On one hand, a citizen science initiative such as eBird provides detailed checklists of all the species a human can see from the ground on a given day. However, the content of these checklists does not necessarily reflect the actual distribution of birds aloft; rather, it favors more conspicuous species, those flying at lower altitudes, those present on the observation site during daytime hours, and those which migrate over more populated areas. This is a form of selection bias, which may be reduced via species distribution modeling but not canceled entirely [13]. Besides, in the context of conservation science, eBird is a powerful tool when applied to species with moderately high abundance but loses in efficiency when monitoring the population of critically endangered species (appearing in almost zero checklists) as well as overabundant species (appearing in almost all checklists).

On the other hand, monitoring bird migration via radar scans leads to an unbiased measurement of the total biomass aloft at the scale of about 1 km^2^. However, this measurement lacks anatomical information which could allow species classification [14]. As a result, when monitoring multiple species of similar body mass yet disparities in conservation status, radar scans alone cannot disambiguate the contribution of each species to the total population in the large-scale migratory flock under study.

These shortcomings motivates the need for a method in addition to weather radar and human observations: the deployment of an acoustic sensor network of autonomous recording units (ARUs) [15–17]. Unlike human observers, an ARU can operate almost anywhere, even the most remote locations, without disturbing the ecological community under study. It can operate 24 hours a day, independently of temperature or sky visibility. Meanwhile, unlike in radar scans, each species of passerine *(Passeriformes)* distinguishes itself from all others by a unique acoustic signature: its flight call.

Flight calls differ from bird songs in terms of their spectrotemporal characteristics: while songs comprise multiple “syllables” and tend to last multiple seconds, a flight call consists of a single acoustic event and lasts between 50 and 150 milliseconds [18]. Such a short time span poses a challenge for identification, be it human or automatic. However, it also presents an opportunity to measure per-species vocal activity not just in terms of presence vs. absence, but in terms of abundance as well (e.g., flight call count over a predefined unit of time). It stems from the above that supplementing human daytime observations and radar scans with acoustic data could, in the near future, improve the reliability of bird migration monitoring; and ultimately evaluate the effectiveness of public policies for avian conservation.

### Computational bioacoustics with deep learning

Despite the growing interest for bioacoustic analysis in avian ecology, the scalability of ARU deployment is currently hampered by the shortage of human experts that are trained to pinpoint and identify bird vocalizations in continuous audio recordings. Indeed, although manufacturing an ARU and collecting audio data is relatively inexpensive, audio annotation is a slow and tedious task [19]. In this context, closing the discrepancy between the cost of hardware and the cost of human labor is crucial to achieving the long-term goal of enabling the deployment of an acoustic sensor network for bird migration monitoring at the continental scale [20].

In order to reduce the annotation overhead involved in bioacoustic monitoring, one promising solution consists in replacing human experts by “machine listening” software. In the ornithological literature, there is a long history of engineering automated or semi-automated bird call detectors, which are initially designed as simple template matching algorithms. These algorithms do not need large computational resources to run, nor large amounts of training data to be parametrized; however, their practical usability is limited to very specific recording conditions, with little or no background noise and a short distance between bird and sensor. In addition, even in the case of a correct detection, they are often unable to distinguish closely related species. As a consequence, human annotation remains the only reliable solution to count bird calls in real-world ARU audio, where the audio signal is polluted by the presence of noise: insects, vehicles, rain, and so forth.

The situation changed recently with the introduction of machine learning, and deep learning in particular, to the field of detection and classification of acoustic scenes and events (DCASE). As the past years have witnessed a relative democratization of high-performance computing (HPC), it has become possible to design more ambitious software architectures for species classification of bird songs and calls. Whereas template matching systems are based on a handful of numerical parameters, deep learning systems typically contain 10^6^ independent parameters or more. These parameters are not tuned by hand, but instead jointly optimized on some pre-annotated training data in order to maximize classification accuracy. The advantage of this increase in dimensionality lies in the robustness of the resulting system: with machine learning, it is theoretically possible to detect bird calls despite high levels of noise and at a long distance. As a downside, machine learning needs a vast amount of training data in order to avoid statistical overfitting—that is, a large gap in classification accuracy between the subset of audio recordings that is annotated and the complementary subset of test data, which may exhibit a different background noise profile. We refer to [21] for a recent review of the state of the art in computational bioacoustics with deep learning.

### Bioacoustic sensor networks as cyberphysical systems

In full generality, a cyberphysical system (CPS) is a network of multiple hardware elements which interact with a complex physical process via a distributed algorithm, thus producing a monitoring tool which may eventually inform decision-making [22]. A well-known example of a CPS is the “smart grid”, an infrastructure in which algorithms monitor and control electrical supply so as to meet constraints of sustainability and reliability. A recent publication has pointed out that the SONYC acoustic sensor network for urban noise monitoring is also a CPS in the sense that machine listening software and microphone hardware constitute its “cyber” and “physical” components respectively [23]. The term CPS has also been employed in the context of acoustic communication between autonomous underwater vehicles [24] and in the SCENA-RBD project for terrestrial monitoring of amphibians [25].

In all the examples listed above, the main challenge facing the deployment of the CPS lies in its multiple spatiotemporal scales of interaction with the data. We point out that bird migration monitoring also fits this definition. From the perspective of bioacoustics, its multiscale temporal extent spans eleven orders of magnitude: the carrier frequency of flight calls belongs to the range 2–10 kHz, hence an oscillation period of the order of 3 · 10^−4^ seconds; whereas the annual migration cycle has a period of 3 10^7^ seconds. In between those two extremes, the time span between two flight calls is equal to 10^2^ seconds, with high variations depending on location and time of day. It thus follows that flight calls make up for about 0.1% of the ARU’s uptime, with the remaining 99.9% being spent in acquiring irrelevant audio data. Likewise, in the spatial domain, North American passerines cover a territory of about 10^13^ square meters yet their flight calls can only be heard over an area of about 10^4^ square meters [18, 26]. The gap between these orders of magnitude implies that bioacoustic sensor networks for bird migration monitoring cannot realistically seek an exhaustive acquisition of all flight calls from Passeriformes [27]. Although they can provide relevant information about species abundance on their own, their ultimate purpose is to serve a complementary source of information to radar aeroecology and crowdsourced observations [28]. Such need for a multiscale interoperability between heterogeneous modalities of data acquisition is what makes bird migration monitoring a prime example of CPS.

However, the design of a CPS necessarily incurs practical risks in terms of reliability, and the case of bioacoustic sensor networks is no exception. Indeed, the ARUs, which constitute the physical frontend of the system, are typically made from low-cost parts [29], deployed in extreme weather conditions [30], and under strong constraints of energy supply [31]. On the “cyber” side of the CPS, machine listening remains a fallible technology, with multiple forms of technical bias. Prior research has shown that the error rate of a state-of-the-art deep learning system for species-agnostic flight call detection may be anywhere between 2% and 20% depending on sensor location; and that its recall may vary between 10% and 70% from dusk to dawn [32]. More generally, audio classification models for bioacoustics should be understood as imperfect approximations of the human ear, and their spatiotemporal predictions of vocal activity as inherently uncertain.

Yet, a search of the existing literature on flight call detection and classification reveals that the approach is rarely posed in its full complexity, but merely as a proof of concept [33–36]. In particular, the machine listening component relies on small–scale data: between 10^2^ and 10^4^ examples in total. Furthermore, these examples tend to originate from field guides and audio recordings in captivity; yet, a prior publication has shown that the classification accuracy of flight calls is lower on field recordings than on captive recordings, across multiple machine listening methods [37]. The number of recording locations and diversity of background noise conditions at training stage thus tends to be insufficient to reflect a practical deployment, in which unforeseen failures necessarily happen [38].

### Contributions

In this article, we present BirdVox, a project for bird migration monitoring via machine listening techniques. Compared to previous studies on flight calls, the main novelty of BirdVox resides in its unprecedented scale: nine sensors which span an area of 10^9^ m^2^ and remain active for 10^2^ days, containing 10^5^–10^6^ flight calls from 10^2^–10^3^ species. The BirdVox project encompasses two contributions: first, a robust machine listening pipeline for the automatic detection and classification of flight calls, named BirdVoxDetect; and secondly, a large-scale evaluation of BirdVoxDetect versus competing methods in real-world conditions of deployment, by way of a new dataset named BirdVox-full-season.

Figure 1 outlines the functioning of BirdVox. The pipeline comprises six stages in total, which are grouped into three blocks: audio signal processing, machine learning, and statistical modeling. The first block involves only engineered transformations and incurs no learning. The second block is data-driven and requires supervised learning on annotated datasets at training time but no further annotation at deployment time. These two blocks form a suite of software tools named “BirdVoxDetect” and “BirdVoxClassify”, for flight call detection and species classification respectively.

**Figure 1.**
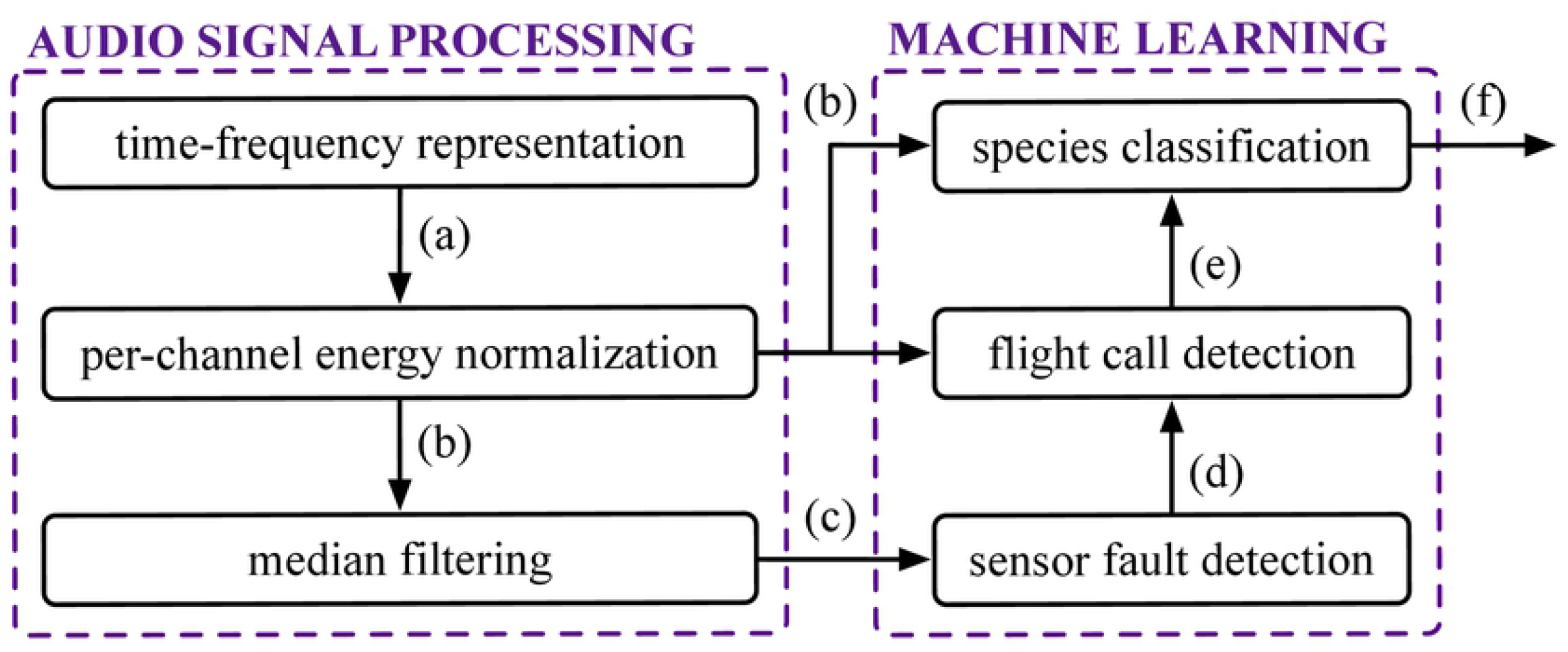
General flowchart of BirdVox, grouped into three blocks. Letter (a) to (f) correspond to subfigures of Figure 2.

Figure 2 presents the output of some key intermediate stages of BirdVoxDetect; i.e., a full night of bird migration from 6 p.m. to 6 a.m. For the sake of clarity, subfigure numbers from (a) to (f) correspond to arrows in Figure 1. Note that, in this case, the sample input presents temporal regions of audible sensor faults, both at dusk and dawn. The BirdVoxDetect pipeline integrates a sensor fault detector which automatically flags these regions in red in visualizations (d) to (f), thus preventing potential false positives. Subfigure 2 (f) presents the result of the second block (“machine learning”) in the BirdVoxDetect pipeline, which is potentially re-usable beyond BirdVox. This subfigure displays flight call timestamps for seven species of passeriformes in addition to a catch-all row of timestamps (in red) for other species.

**Figure 2.**
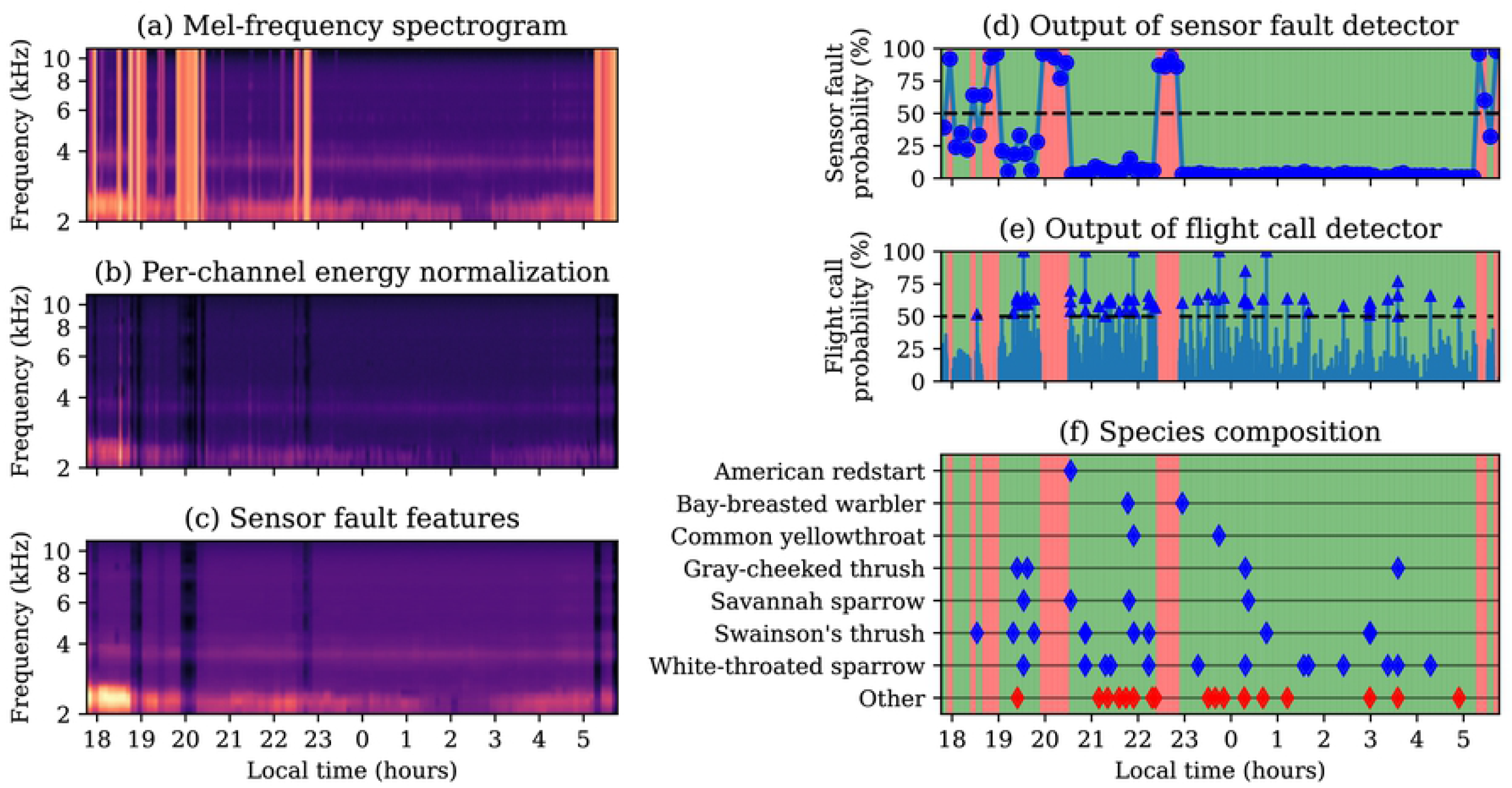
Sample output of BirdVox. Brighter colors in subfigures (a) to (c) denote larger values in the time–frequency domain. Red regions in subfigures (d) to (f) denote time segments where a sensor fault is detected. Every triangle in subfigure (e) represents a flight call, as Every blue lozenge in subfigure (f) represents a flight call from an identifiable species.

The second contribution of our paper is the open release of the largest audio dataset of flight calls to date, named BirdVox-full-season (or full-season for short)^1^. The full-season dataset contains 6671 hours of audio across nine sensors from Tompkins County, NY. Furthermore, an expert (AF of the authors) has annotated two of its subsets: BirdVox-full-season (or full-season for short) and BirdVox-296h (or 296h for short). The former contains 62 hours of continuous audio from one night within the full-season and serves for training of the flight call detector in BirdVoxDetect. The latter contains 150 two-hour audio segments which span the spatiotemporal and acoustical diversity of the full-season.

Figure 3 lists all the audio datasets of the BirdVox project. Note that, because BirdVox-296h is disjoint from the training of BirdVoxDetect, thus allowing us to use it as an evaluation benchmark. Once the evaluation is complete, we deploy BirdVoxDetect on the full-season dataset via a massively parallel computing task comprising around 1.6 · 10^9^ Fourier transforms and 4.8 · 10^8^ neural network predictions.

**Figure 3.**
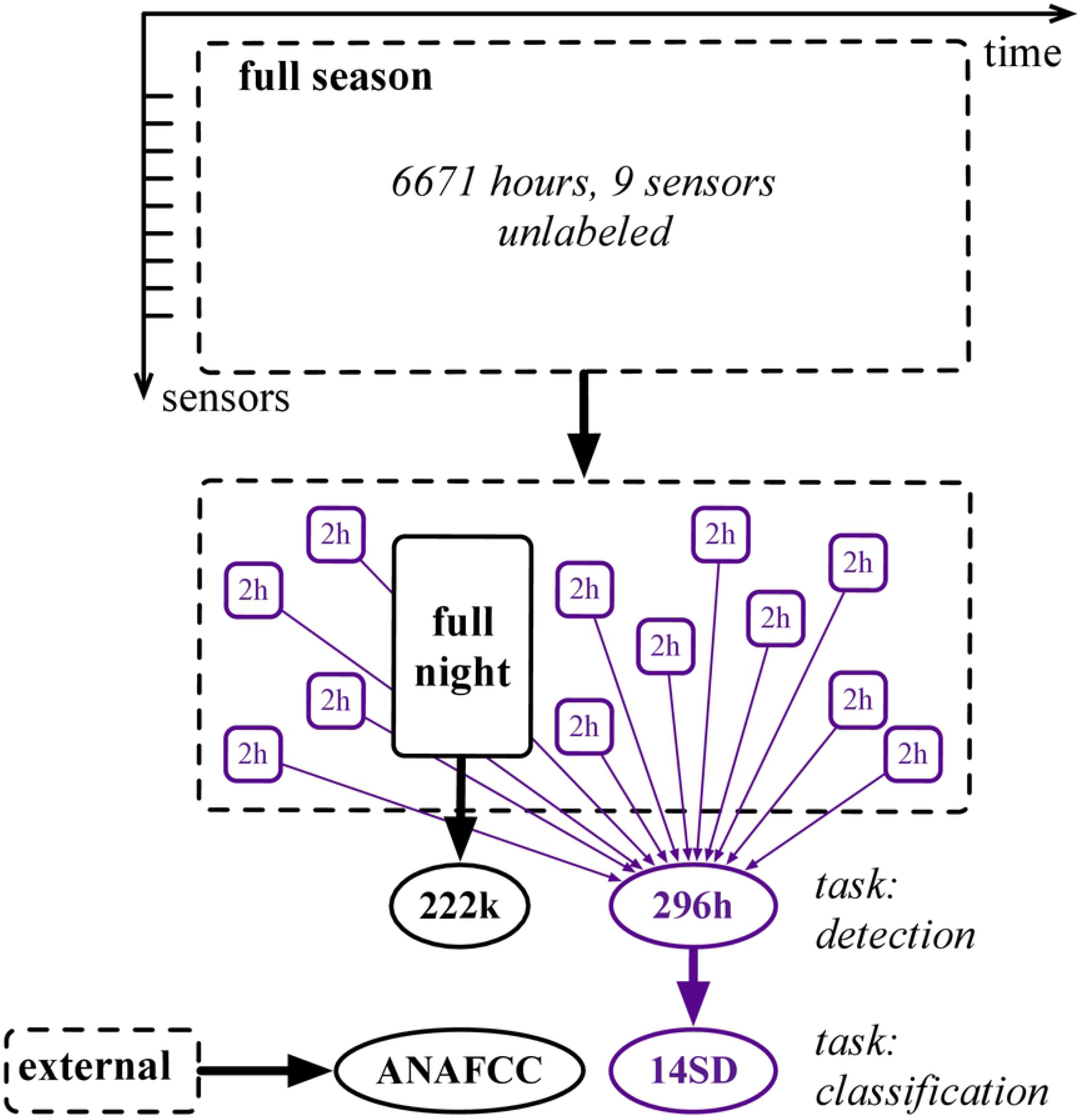
Diagram of the full-season dataset and its subsets. Solid and dashed lines denote labeled and unlabeled audio respectively. Black and purple lines denote training and evaluation subsets respectively. Rectangles and ellipses denote full-length acoustic scenes and isolated audio clips respectively.

## Data collection

This section presents our procedure of audio acquisition and expert annotation of acoustic events.

### Deployment of a bioacoustic sensor network

In 2015, we placed nine bioacoustic sensors in residential areas surrounding the town of Ithaca, NY, USA. All sensors in our deployment setting correspond to the same hardware specification: namely, the Recording and Observing Bird Identification Node (ROBIN) developed by the Cornell Lab of Ornithology. Each ROBIN comprises a Knowles EK23132 microphone element, an analog-to-digital converter, a Raspberry Pi Model B single-board computer, a solid-state memory card, and a battery. The microphone element is omnidirectional and has an approximately flat sensitivity of 53 ± 5 dB between 2 and 10 kHz; that is, the frequency range of flight calls [39]. The microphone element sits at the bottom of a small horn-shaped enclosure oriented upwards. In turn, this enclosure sits inside a hard plastic housing, whose purpose is to reject lateral sound sources, such as insects or car engines^2^.

The analog-to-digital converter encodes the monophonic signal recorded by the microphone into a linear pulse-code modulation sequence at a sample rate of 24 kHz and a sample depth of 16 bits. The single-board computer streams this sequence under the form of 20-second buffers, which are progressively appended to a lossless audio file in FLAC format. This acquisition procedure is repeated every night from dawn to dusk between August 3^rd^, 2015 and December 8^th^, 2015. This corresponds to roughly 1,500 hours of audio per sensor, and thus 13, 500 hours for the entire sensor network. However, due to intermittent failures of sensing hardware, only 6, 671 hours were successfully retrieved.

Figure 4 presents the spatial distribution of sensors in Tompkins County, NY, USA. We observe that the availability of audio data varies starkly across sensor locations between 107 and 1, 356 hours, with a median of 834 hours. Furthermore, the sensor network does not follow a simple geographical pattern, such as a uniform linear array or a rectangular grid. Indeed, for reasons of privacy and practicality, the sensors were deployed in backyards or on house roofs, pertaining to members of the Cornell Lab of Ornithology (see Acknowledgments section).

**Figure 4.**
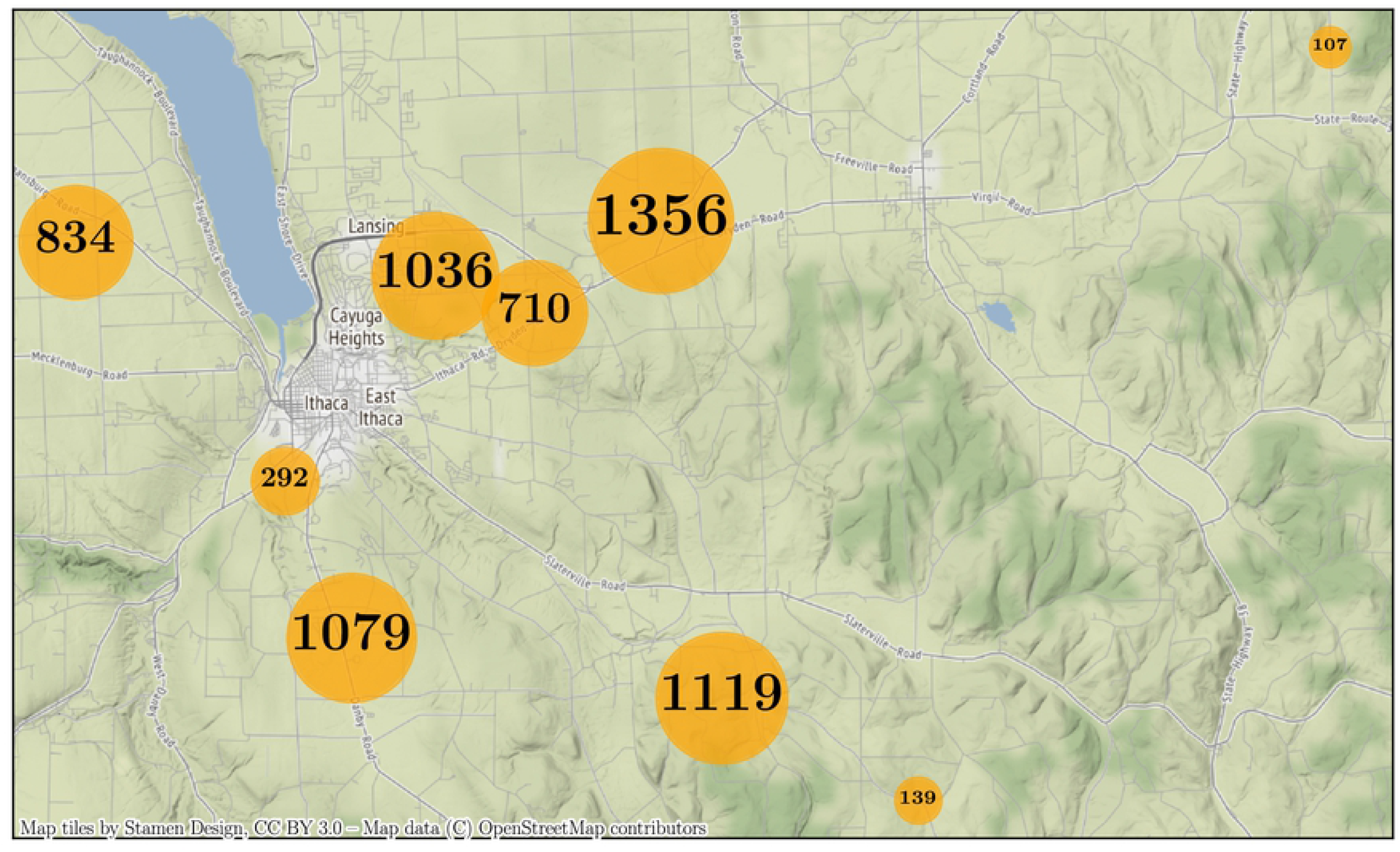
Map of sensors in the full-season dataset. The map shows the surroundings of Ithaca, NY, USA, over an area of roughly 1.000 km^2^, i.e. 40 km from West to East and 25 km from North to South. The area of each orange dot is proportional to the total duration of available audio in the corresponding sensor.

### Expert annotation of flight calls

We divide all recordings in the full-season dataset into two-hour segments. The starting times of these segments are expressed in Coordinated Universal Time (UTC) and range from 6 p.m. to 6 a.m. by increments of two hours. Note that the local time in Ithaca, NY, corresponds to Eastern Standard Time (UTC-05:00) in winter and Eastern Daylight Time (UTC-04:00) in summer. Furthermore, for each nocturnal recording in full-season, we extract the audio segment corresponding to the two hours preceding sunrise. To determine the time of sunrise on any given day, we rely on open data from the weather station of the Ithaca Tompkins Regional Airport (KITH). This operation results in a collection of 3, 131 segments, amounting to 6, 262 hours of audio.

We select 150 segments at random from the aforementioned collection. Among them, 100 segments are synchronized with UTC, ranging between 6 p.m. and 6 a.m, while the remaining 50 correspond to the two hours preceding sunrise. The reason why we assign a larger relative proportion to the latter is that the density of flight calls is higher at dawn than at dusk or at night [40].

In 2018 and 2019, an expert ornithologist (AF of the authors) annotated each of these 150 segments by means of the Raven Pro sound analysis software^3^. The annotation task consisted in pinpointing and labeling every flight call in the time–frequency domain. It took 570 hours to complete this first round of annotation. A second round of annotation, conducted in 2021, revealed that two segments were not admissible for nocturnal flight call detection because they had mistakenly been extracted after sunrise. After excluding these two segments, we obtained 148 segments, corresponding to 296 hours of audio.

The annotation files for those 296 hours comprise over 100 distinct sound categories. We filter out categories corresponding to non-animal sounds (e.g., *alarm, rain*), invertebrate sounds (*katydid*), non-bird sounds (*frog, coyote*), and non-passeriforme bird sounds (*Caspian Tern, Green Heron*). Then, we focus on a list of 14 birds of interest: four American sparrows, one cardinal, two thrushes, and seven New World warblers (see Figure 5). Outside of these four families, we aggregate all flight calls from Passeriformes under a common catch-all category: “other Passeriformes”, e.g., *American Goldfinch, Baltimore Oriole, Golden-crowned Kinglet*. Furthermore, we build catch-all categories for each of the four passerine families of interest. For example, “other American Sparrows” includes eight species, e.g. *Field Sparrow*. Likewise, “other Cardinals” includes three species, e.g. *Indigo Bunting*; “other Thrushes” includes four species, e.g. *Veery*; and “other New World Warblers” includes 16 species, e.g. *Magnolia Warbler*.

**Figure 5.**
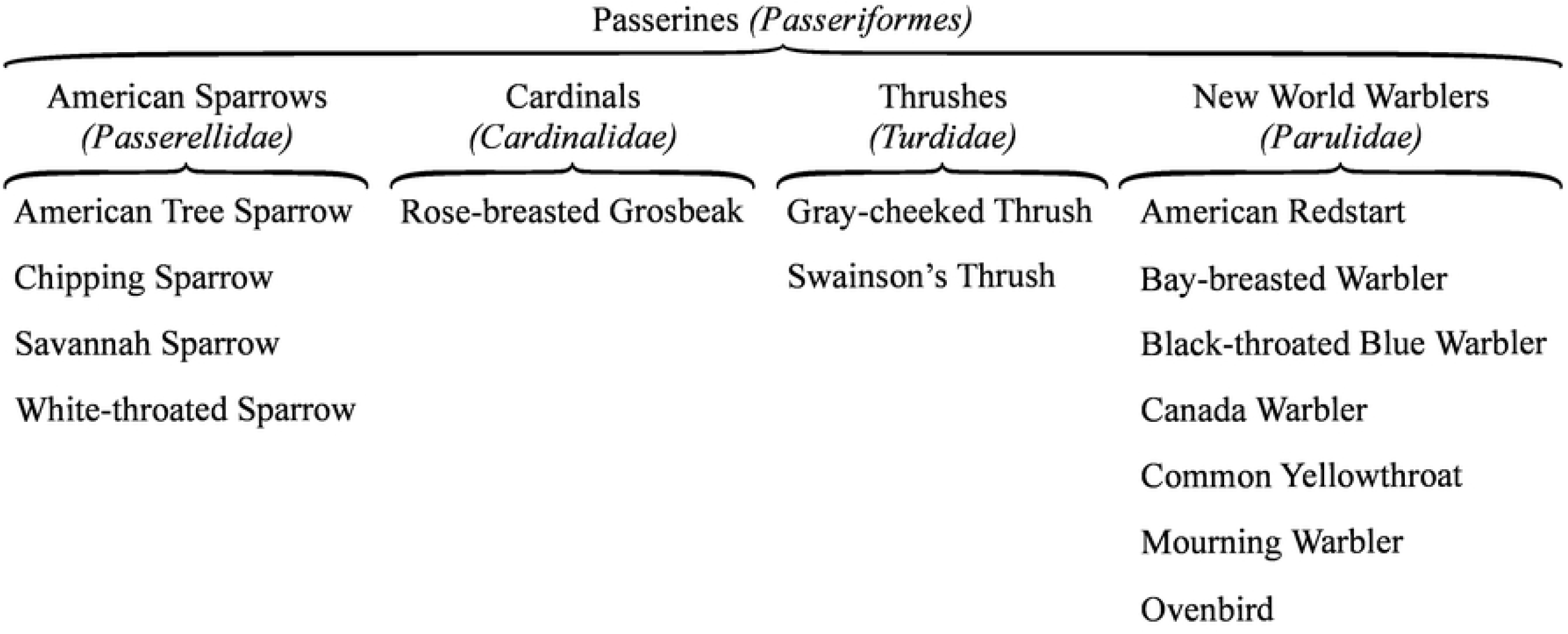
Taxonomy of labels in the 296h dataset. The coarse, medium, and fine level of the taxonomy correspond to order, family, and species respectively. Species within the same bracket belong to the same family of the *Passeriformes* order.

Due to the varying distance between the sensor and the source, some of the flight calls are too faint to be confidently labeled in terms of species, even to an expert ear. However, they may be identifiable at a coarser taxonomic level. In those instances, automatic species classifiers can only be evaluated against the human ground truth up to a certain level of granularity [41]. For this reason, we release three variants of the annotation, respectively denoting order, family, and species.

We name BirdVox-296h (or “296h” for short) the annotated dataset of 150 two-hour segments, as a subset of BirdVox-full-season. For the sake of research reproducibility, we upload a copy of BirdVox-296h to the Zenodo repository of open-access data^4^.

### Per-channel energy normalization (PCEN)

Let **E**(*t, f*) be the mel-frequency spectrogram of some audio recording, with *t* and *f* denoting discrete time and mel frequency respectively. We define a low-pass filter *φ* _*T*_ with a cutoff frequency equal to *T* ^−1^. Per-channel energy normalization (PCEN), originally introduced by [42], applies adaptive gain control and dynamic range compression **E** by means of the following equation:

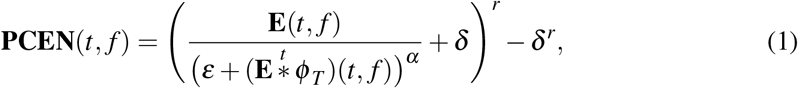

where the quantities *ε, α, δ*, and *r* are constants and the notation 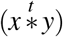 denotes a convolution over the time dimension. In practice, we construct *φ* _*T*_ as a first-order IIR filter whose response to **E** is of the form

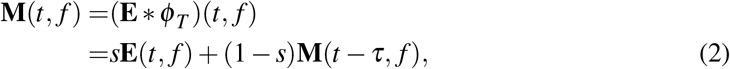

where the constant *s* is the weight of the associated autoregressive process (AR(1)) and *τ* = 1.5 ms is the hop size of the mel-frequency spectrogram. The recursive implementation above is more computationally efficient than FFT-based convolution while having a smaller memory footprint. Proposition IV.1 from [43] gives the following connection between constants *T, τ*, and *s*:

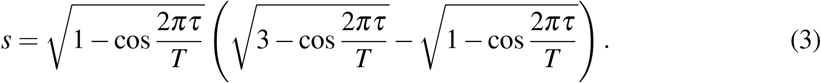

In this paper, we set *T* = 60 ms, *ε* = 10^−6^, *α* = 0.8, *δ* = 10, and *r* = 0.25. A previous publication [32, Figure 4 specifically] has shown empirically that, on the full-night dataset, this choice of parameters Gaussianizes the distributions of normalized spectrogram magnitudes, consistently across sensors. This is in contrast with the original publication on PCEN [42], in which the proposed default parameters are optimized for automatic speech recognition in a noisy indoor environment, but inadequate for flight call detection.

In the rest of this paper, we refer to the output of PCEN by the abbreviation “PCEN-gram”. Figure 2b illustrates a sample PCEN-gram.

### Median filtering

We compute the running median of the PCEN-gram over non-overlapping windows of duration 30 minutes, for each mel-frequency subband *f* independently. This results in a time–frequency representation in which the time axis is sampled at a rate of two frames per hour. Furthermore, we subsample the mel–frequency axis by a factor of 12, thus reducing the number of subbands *f* from 120 down to 10. We call “sensor fault features” the resulting time–frequency representation, samples at a rate of two frames per hour. Figure 2c illustrates a sample output of median filtering.

### Sensor fault detection

This section presents the sensor fault detector of our system, i.e., a random forest classifier trained with an active learning paradigm. In the functional diagram of Figure 1, the sensor fault detector corresponds to block (d).

### Random forest classifier

We extract sensor fault features on the full-season dataset. This operation results in 12k feature vectors, i.e., one every half-hour segment. We manually label two half-hour segments in the dataset: one in which a sensor fault is present and the other in which no sensor fault is present. With scikit-learn v0.20.1 [44], we train a random forest classifier on the two sensor fault feature vectors corresponding to these two segments. We set the number of ensembled decision trees in the random forest equal to 100.

### Active learning for efficient audio annotation

Because the classifier described above is trained on a tiny dataset (two samples), it does not generalize well to unseen recording conditions. To improve accuracy, it is necessary to refine the decision boundary between classes, and thus label more samples. However, the annotation of sensor faults from bioacoustic recordings is a particularly tedious task. Furthermore, the relatively rare proportion of sensor faults in full-season (estimated between 1% and 5% of the audio data) causes a class imbalance problem, which hampers the statistical generalization of the classifier.

We address the issue of annotation efficiency in the sensor fault detection task by adopting an active learning paradigm. Instead of annotating audio segments drawn uniformly at random in full-season, we execute an algorithm which iteratively queries the human annotator with the most informative unlabeled sample. Here, the informativeness of a sample is defined according to the prediction confidence of the random forest classifier.

We apply the active learning algorithm of [45], known as “alternate confidence sampling”. In *f*_HC_ = 90% of the iterations, the algorithm queries the human with the unlabeled sample with least confidence, that is, the one closer to the decision boundary of the classifier. Alternatively, in one every ten iteration, the algorithm queries the human with a high-confidence sample: specifically, one sample drawn uniformly at random among the pool of unlabeled samples whose confidence exceeds a fixed probability threshold of *T*_HC_ = 85%.

The human annotator labels samples progressively, as queried by the active learning algorithm. Conversely, the random forest classifier is retrained after the labeling of every sample, and thus becomes progressively more discriminative. This human-in-the-loop machine learning procedure is repeated until the classifier reaches a satisfying generalization power. In practice, two annotators (AF and VL of the authors) labeled 100 half-hour segments in full-season.

Figure 6 illustrates the prediction of the sensor fault classifier on the entire full-season dataset, after being trained on these 100 half-hour segments. We observe that sensor faults (in orange) affect all eight of the nine sensors intermittently and tend to span across two or more consecutive nights.

**Figure 6.**
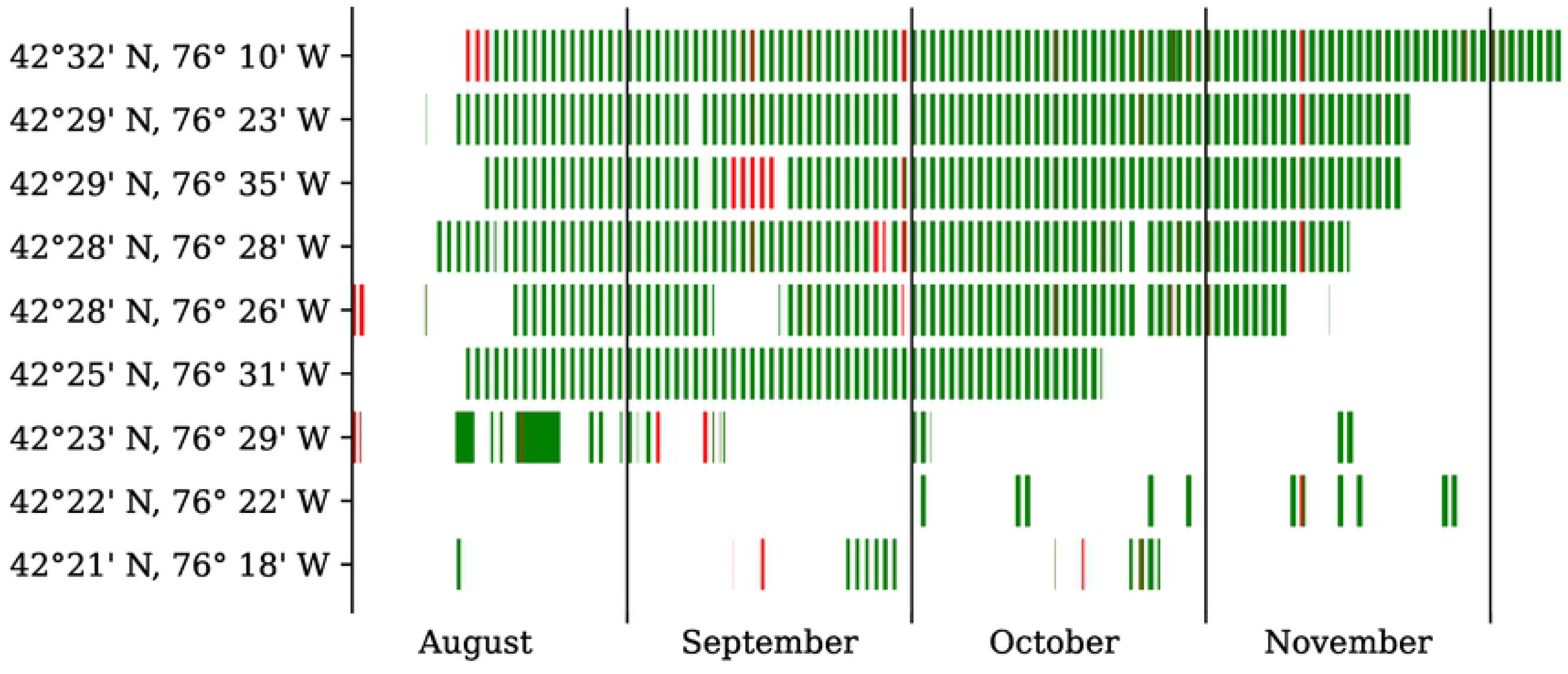
Calendar of recordings in the full-season dataset, organized by month (x-axis) and by uptime (y-axis). Every red (resp. green) rectangle represents a faulty (resp. non-faulty) recording, as determined by our random forest classifier.

### Qualitative evaluation with *t*-SNE embedding

We propose to shed light on the active learning process described above by visualizing the performing a *t*-distributed stochastic neighbor embedding (*t*-SNE) of the full-season dataset [46]. The *t*-SNE algorithm learns a nonlinear mapping from a feature space in dimension ten to a embedding space in dimension two. In doing so, *t*-SNE minimizes the Kullback-Leibler divergence between the joint probability distribution of samples in the feature space and that of samples in the embedding space. Therefore, spatial proximity in the 2-D embedding space denotes acoustical similarity in terms of median PCEN-gram features. We use the implementation of scikit-learn with all parameters set to their default values as of v0.20.1.

Figure 7 illustrates the outcome of *t*-SNE embedding. In the left column, we represent unlabeled samples as black dots and labeled samples in color: green square for positives (i.e., absence of sensor fault) and red squares for negatives (i.e., presence of sensor fault). In the right column, we represent the predictions of the sensor fault detector over all samples, be them labeled or unlabeled: darker shades of red (vs. green) denote a greater predicted probability that a sensor fault is present (resp. absent) in the corresponding audio excerpt. We repeat the display at different stages of the active learning process: initialization (top), with 10 labeled samples (center), and with 100 labeled samples (bottom).

**Figure 7.**
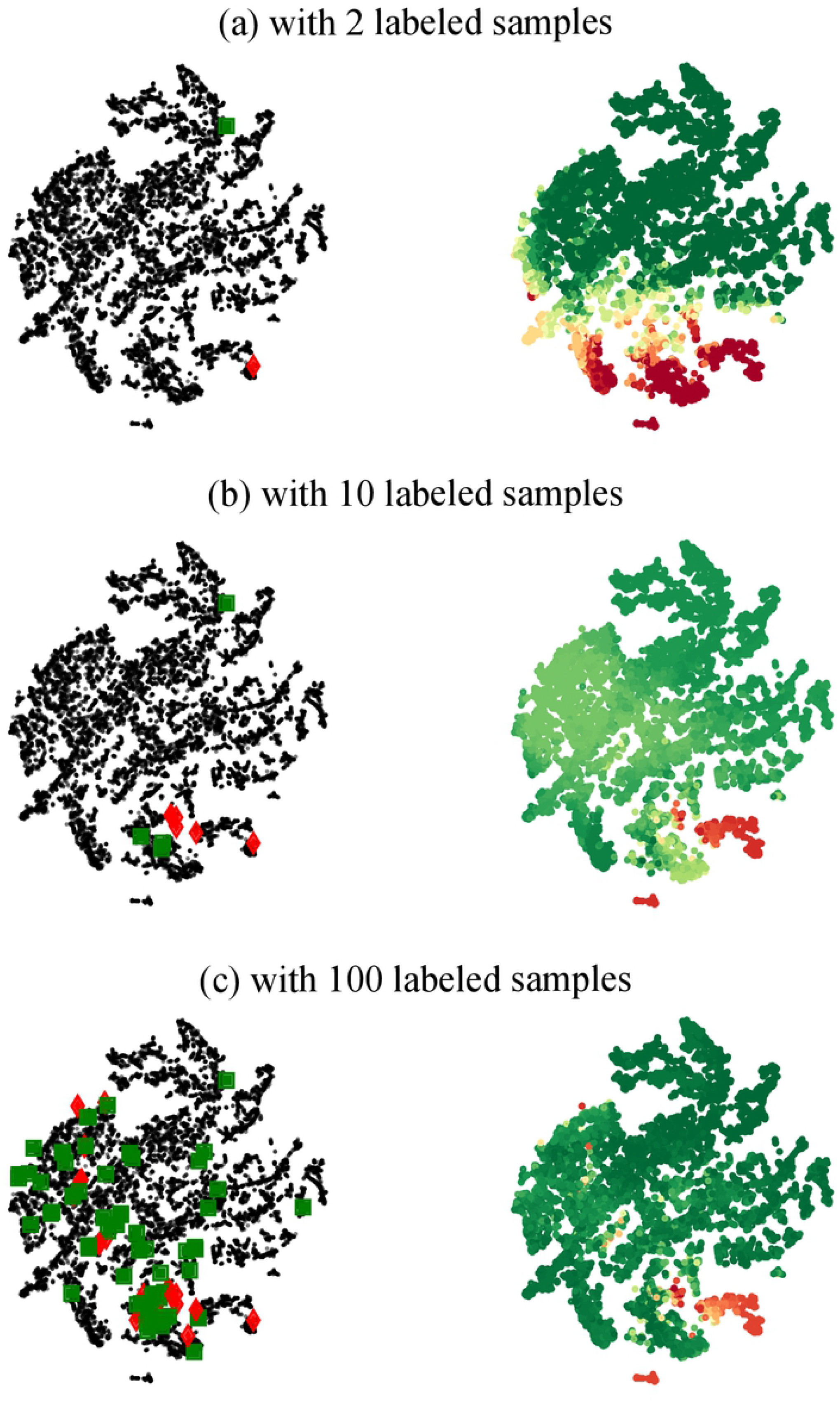
Visualization of sensor fault features with *t*-SNE. Left column: human expert annotation. Red lozenges (resp. green squares) denote the presence (resp. absence) of a sensor fault in the corresponding audio excerpt. Unlabeled samples are denoted as small black dots. Right column: machine listening prediction by a random forest classifier trained on sensor fault features in dimension ten. Darker shades of red (vs. green) denote a greater predicted probability that a sensor fault is present (resp. absent) in the corresponding audio excerpt.

We observe in Figure 7c (left) that the distribution of labeled samples is not uniform over the *t*-SNE map. Instead, it is concentrated on the regions of least confidence of the sensor fault detector: the top-left and bottom-right corners of the scatter plot in our case, appearing in pale green in Figure 7b (right). Moreover, we observe on Figure 7a (right) that the decision boundary of the sensor fault detector appears as a rectilinear color gradient at the initialization. In contrast, we observe on Figures 7b (right) and 7c (right) that the decision boundary becomes progressively sharper and nonlinear as the number of labeled samples increases. These observations provide qualitative evidence that the proposed active learning process accelerates the convergence of the sensor fault detector as a function of training set size.

### Flight call detection

This section presents our deep learning system for species-agnostic avian flight call detection, named BirdVoxDetect. BirdVoxDetect is a convolutional neural network (CNN) taking a PCEN-gram representation as its input. In the functional diagram of Figure 1, BirdVoxDetect corresponds to block (e).

### Deep learning architecture

BirdVoxDetect draws its inspiration from prior research on urban sound classification [47] and species classification from clips of flight calls [48]. It composes three convolutional layers and two fully connected layers. The first (resp. second) convolutional layer consists of 24 (resp. kernels of size 5 × 5, a rectified linear unit (ReLU) activation function, and a strided max-pooling operator of shape 4 × 2; that is, 4 time frames and 2 frequency bands. The third convolutional layer consists of 48 kernels of size 5 × 5, a ReLU, and no pooling. The first fully connected layer contains 64 hidden units, followed by a ReLU. Lastly, the second fully connected layer maps those 64 hidden units to a single output unit, followed by a sigmoid nonlinearity. Figure 8 illustrates the architecture of BirdVoxDetect, as rendered by the NN-SVG online tool [49].

**Figure 8.**
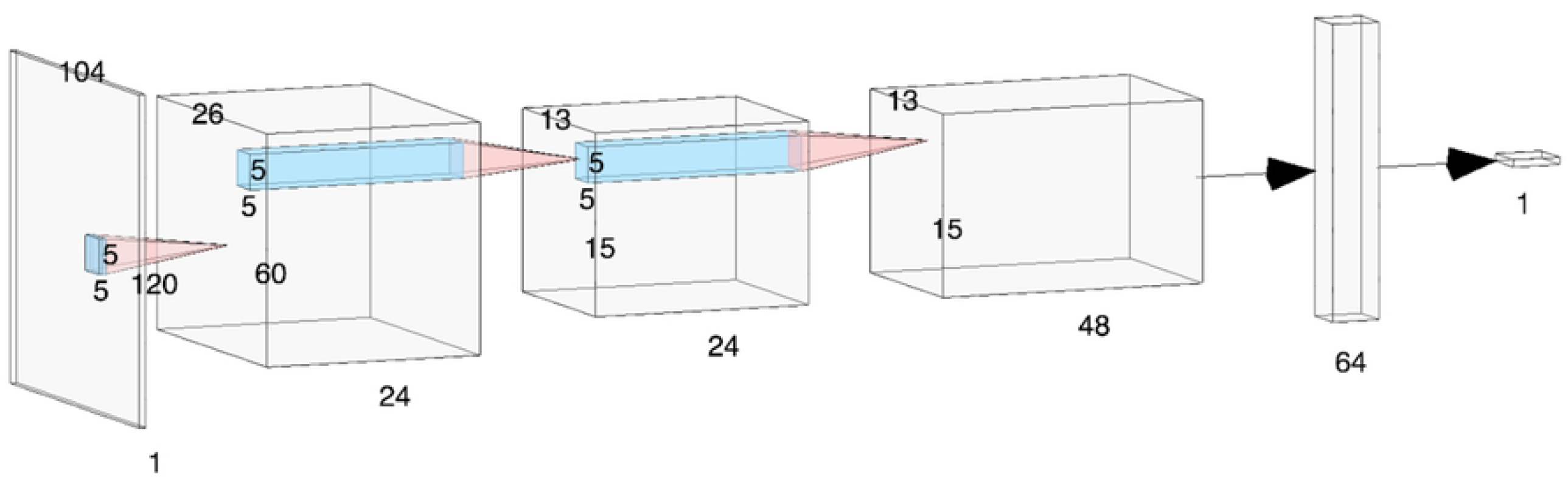
Functional diagram of the convolutional neural network for flight call detection (BirdVoxDetect). Grey tensors represent intermediate computations and blue regions represent receptive fields of convolutional layers.

**Figure 9.**
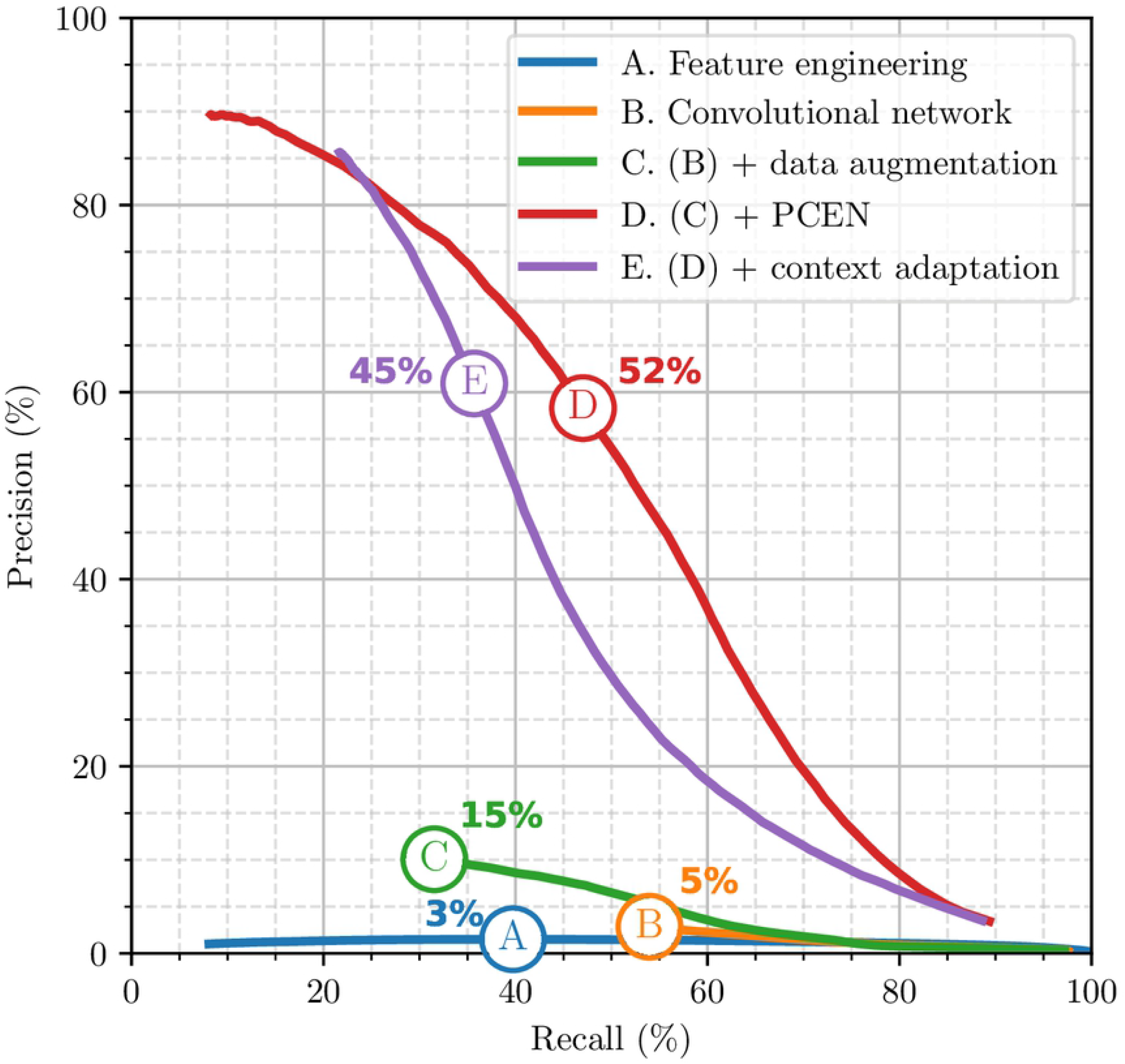
Precision–recall curves of BirdVoxDetect for the 296h dataset. Each dot on the curve denotes a different value of the BirdVoxDetect threshold.

**Figure 10.**
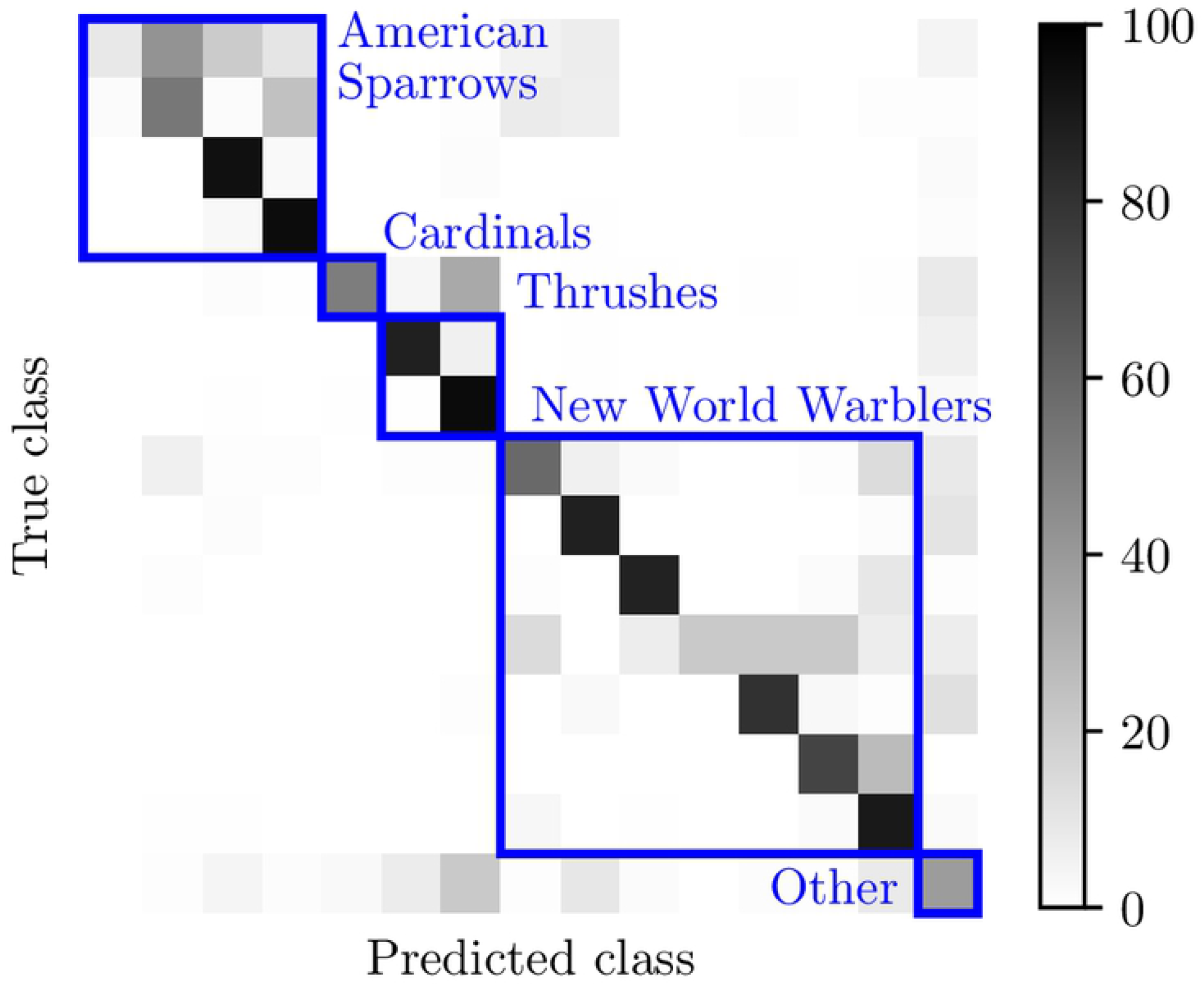
Confusion matrix of BirdVoxDetect on the 296h dataset. The color of each element indicates the percentage points of ground-truth positive examples that are predicted as each class. The species are ordered (downward and rightward) and grouped (in blue boxes) according to the taxonomy in Figure 5.

The input to BirdVoxDetect is a PCEN-gram excerpt of duration equal to 150.9 ms, corresponding to the center portion of the audio clips in BirdVox-222k. This PCEN-gram is encoded as a single-precision real matrix with 120 rows and 104 columns. During training, we apply batch normalization to this matrix (but not to deeper layers), thus bringing its coefficients to null mean and unit variance.

Throughout the development of BirdVoxDetect, we have explored over 100 common variations in hyperparameters: kernel size, layer width, number of layers, mel scale discretization, multiresolution input, choice of nonlinearity, use of dropout, use of batch normalization, and choice of learning rate schedule. Yet, none of these variations improved validation accuracy systematically (see next subsection for details on evaluation). Of course, we do not claim that the architecture presented in the paragraph above is optimal; but our failure to improve it via common hyperparameter variations does suggest that it faithfully represents the accuracy of CNNs—in the sense of a general family of machine learning models—on the flight call detection task.

### Data curation

We train BirdVoxDetect on BirdVox-222k (or “222k” for short), a new derivative of the BirdVox-full-night dataset. BirdVox-full-night (or “full-night” for short) comprises 62 hours of audio in total, as recorded on the night of September 23rd, 2017 by six different sensors. In 2017, an expert ornithologist (AF of the authors) spent 102 hours annotating each of these six recordings and found 35k flight calls from passeriformes.

We extract 35k audio clips from full-night, each lasting two seconds and centered around one annotated flight call. We group these 35k audio clips into 352 segments, each of them of size 100, according to their spatiotemporal contiguity in the sensor network. Then, we run a pretrained flight call detector full-night: this detector combines spherical *k*-means (SKM) and a support vector machine (SVM), and is thus an instance of “shallow learning”. We use the false alarms of this detector as a source of challenging negatives for the spatiotemporal region corresponding to each segment. By design, the negative-to-positive ratio varies between 1 and 9 depending on the segment, but is always integer. We refer to [40] for more details on full-night and the shallow learning classifier.

Furthermore, we count the spatiotemporal temporal distribution of flight calls per sensor location and per two-hour segment within the full night. Following this coarse spatiotemporal estimate, we extract 222k audio clips at random within the time regions containing no flight calls. Combining the 35k positive clips (centered around one flight call) and the 187k negative clips (containing no flight call) yields the 222k dataset. Note that the ratio of positives to negatives in 222k is equal to 187k/35k ≈ 5.3.

We divide 222k into a training set and a validation set, following a 85% / 15% random partition. Contrary to prior research on full-night, we do not perform “leave-one-sensor-out” cross-validation but a simple shuffle split without regard for sensor location. Indeed, in this article, we are not primarily interested in the generalization ability of BirdVoxDetect from one sensor to another but from one night of audio acquisition (full-night) to several months (full-season). The training subset of 222k amounts to 299 segments or 189k samples.

### Data augmentation

We augment the 222k dataset with three kinds of digital audio effects: pitch shifting, time stretching, and the combination of pitch shifting and time stretching. We sample the pitch interval at random from a normal distribution with null mean and half-unit variance, as measured in semitones according to the 12-tone equal temperament. Furthermore, we sample the time stretching factor at random from a log-normal distribution with parameters *µ* = 0 and *σ* = 0.05.

Such a randomization procedure allows to augment any given audio sample more than once. Specifically, we draw ten instances of pitch shifting, ten instances of time stretching, and ten instances of pitch shifting and time streching in combination. This corresponds to 31 versions of each audio sample in total: i.e., one original version and 30 augmentations. After augmenting each of the 189k samples in the training set of BirdVox-222k, we obtain a dataset of 31 × 189k = 5.9M samples. This dataset represents 633 gigabytes of data on disk.

### Training

We train BirdVoxDetect on the augmented training subset of BirdVox-222k via the Adam algorithm, an improved variant of stochastic gradient descent. We leave the hyperparameters of Adam to their default values: i.e., a learning rate of 10^−3^, decay rates of *β*_1_ = 0.9 and *β*_2_ 0.999, and a denominator offset of 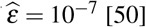 = 10^−7^ [50].

We formulate the flight call detection task as binary classification and choose binary cross-entropy as objective function. Similarly to [40], we regularize this objective function by penalizing the *L*^2^ norm of the synaptic weights in the penultimate layer, with a multiplicative factor set to 10^−3^.

To implement the training procedure efficiently, we use the pescador Python package, which offers utility functions for shuffling and streaming heterogeneous data^5^. For each of the 299 segments and the 31 augmentations, we construct a “stream”: that is, an infinite generator which yields positive and negative samples with equal probability. At the beginning of each epoch, pescador draws one augmentation uniformly at random (out of 31) for each of the 299 segments. We then define batches by “multiplexing” the streams corresponding to these 299 segment–augmentation pairs, so that each stream contributes one and only one sample per batch. In this way, we guarantee that the 299 samples in each batch reflect the acoustical diversity of the full-night dataset. We repeat the process 100 times per epoch, thus yielding 29.9k samples per epoch in total. Note that this number roughly corresponds to the number of flight calls in the training set.

On every epoch, we re-draw a new augmentation for each of the 299 available segments and multiplex the corresponding streamers. Thus, different epochs contain the same original audio material but vary stochastically in terms of augmentations. Furthermore, we guarantee that the spatiotemporal density of negatives matches that of positives. We run Adam for 24 hours on a CPU and checkpoint the model with lowest validation loss.

### Evaluation

After training on 222k, we evaluate BirdVoxDetect on 296h. Note that 222k and 296h arise from the same recording locations but are disjoint in time. Furthermore, the dataset 222k was constructed from a single night of data acquisition whereas 296h is more diverse, as it involves recordings between August and November 2015.

We run BirdVoxDetect on each of the two-hour segments in 296h according to a hop duration of 50 ms, thus producing an event detection function at a rate of 20 Hz. We select local peaks in the detection function above some fixed absolute threshold *τ* ∈]0, 1[. Then, we compare the set of detected peaks to the human-provided checklist of flight call timestamps.

We define matching pairs between detected events and a reference events if their timestamps are within 500 ms of each other. We optimize the cardinality of this matching while guaranteeing that each reference peak matches a single detected peak at most, and vice versa. For this purpose, we solve a bipartite graph matching problem via the match events function of the mir eval Python package [51]. This operation yields a number of true positives, false positives, and false negatives. We may convert these integer counts into information retrieval metrics: precision, recall, and *F*_1_-score. We repeat the process for sweeping values of the threshold parameter *τ* in to derive a precision–recall curve.

## Results

Figure 11 summarizes out results. First, we evaluate a flight call detection system that does not rely on deep learning, but purely on feature engineering. Under the names of “Tseep” and “Thrush”, this system has long remained the standard for detecting sparrows, warblers and thrushes respectively [**?**]. We re-implement these detectors in Python, with help from the original authors. We obtain an *F*_1_-score of 3%. As shown on Figure 11 (curve A), this low *F*_1_-score can be explained by a low precision; that is, a large proportion of false positives in comparison with true positives.

**Figure 11.**
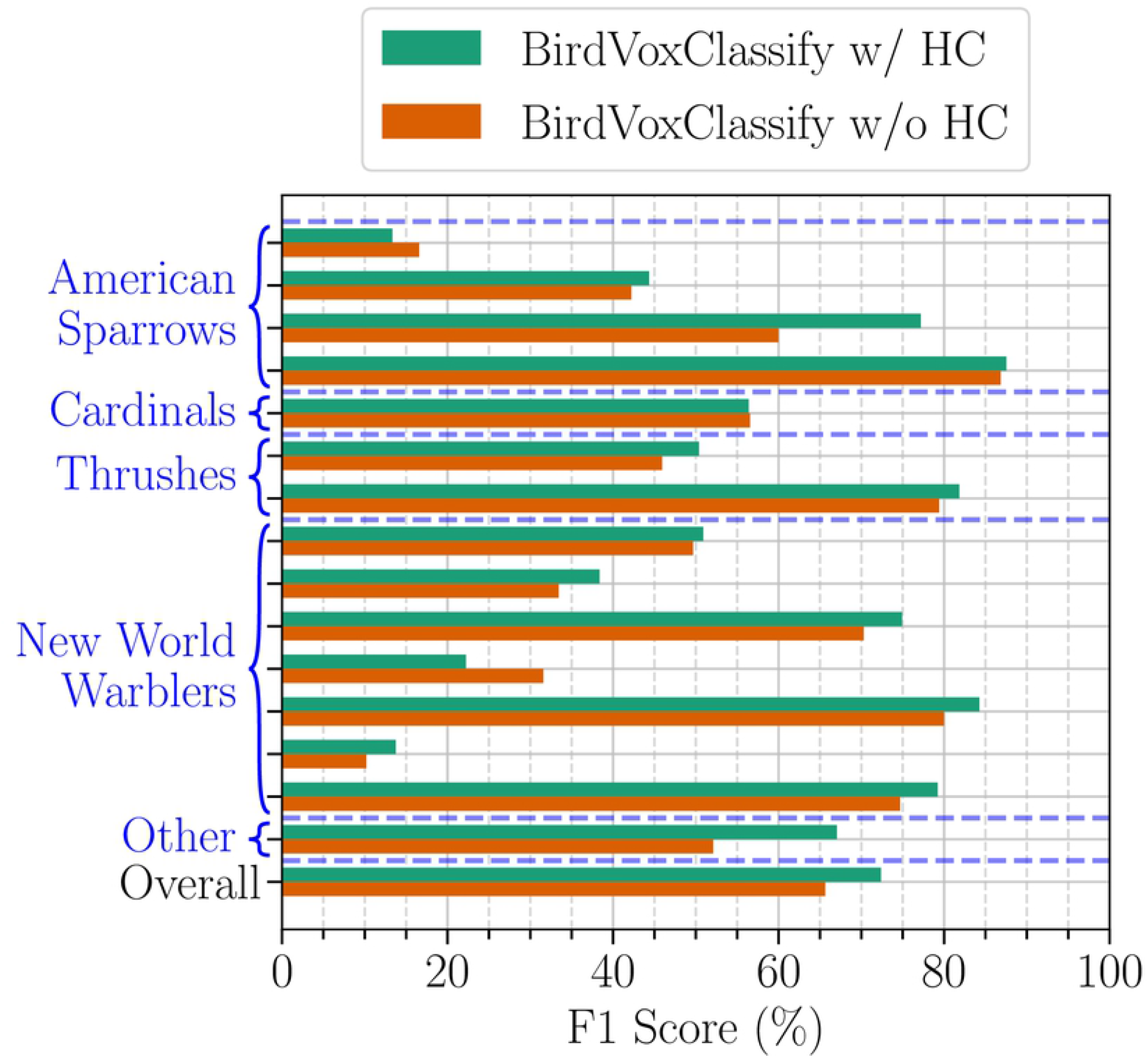
Evaluation of hierarchical consistency on the 296h dataset. Species-specific *F*_1_-scores are ordered (downward) and grouped (in blue brackets) according to the taxonomy in Figure 5. The row “other” represents the *F*_1_-score of out-of-vocabulary samples while the row “overall” corresponds to a micro-averaged *F*_1_-score across all training samples in the dataset.

**Figure 12.**
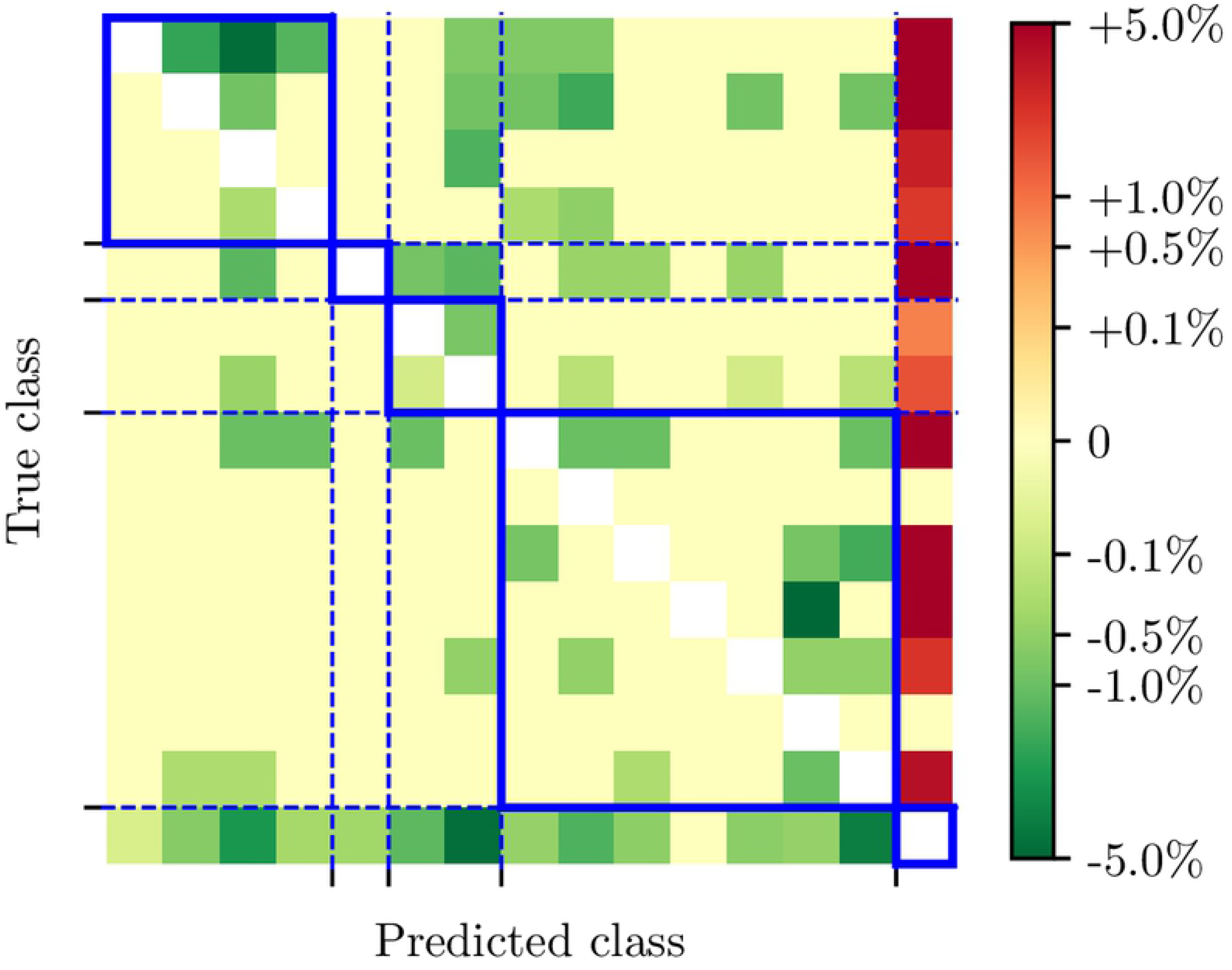
Effect of hierarchical consistency on the confusion matrix. Relative change in class-wise confusion of BirdVoxDetect at the species level for the 296h dataset, without and with hierarchical consistency. Green (resp. red) cells denote a relative decrease (resp. increase) in confusion with respect to the confusion matrix in Figure 11. The species are ordered (downward and rightward) and grouped (in blue boxes) according to the taxonomy in Figure 5.

Then, we evaluate a version of BirdVoxDetect yet without PCEN nor data augmentation. Instead, we train the network on batch-normalized log-mel-spectrograms from BirdVox-222k. We obtain an *F*_1_-score of 5%: see curve B in Figure 11. Artificial data augmentation, as described above, brings the *F*_1_-score of the deep convolutional network to 15%: see curve C. Replacing the log-mel-spectrogram representation by a PCEN-gram considerably improves *F*_1_-score up to 52%: see curve D. Lastly, implementing a form of context adaptation in the neural network, as proposed in [43], reduces the *F*_1_-score to 45%: see curve E. Therefore, we keep model D as our flight call detector of choice and release it publicly under the name of BirdVoxDetect v0.6.

### Species classification

This section presents our deep learning system for multilevel taxonomic avian flight call classification, named BirdVoxClassify. Similarly to BirdVoxDetect, BirdVoxClassify is a CNN taking a 120-band PCEN-gram as its input. Because BirdVoxDetect and BirdVoxClassify share a common input representation, we may pass positive clips from BirdVoxDetect to BirdVoxClassify directly in the PCEN-gram domain instead of the waveform domain, without having to recompute PCEN. In the functional diagram of Figure 1, BirdVoxClassify corresponds to block (f).

### Deep learning architecture

The architecture of our multilevel taxonomic classifier corresponds to a non-hierarcical multitask model (abbreviated Non-H. MT) presented in prior species classification research [52]. Although this prior publication reported that a hierarchically structured classifier (TaxoNet) achieved the best classification performance on its evaluation dataset, we were not able to replicate the results with the new data and now find that the non-hierarchical multitask model performs best.

The architecture of BirdVoxClassify is similar to that of BirdVoxDetect, as it also composes three convolutional layers and two fully connected layers, with no bias weights for any layer. Before the first layer, we perform batch normalization on the PCEN-gram to stabilize and accelerate training [53]. The three convolutional layers are identical in shape to those of BirdVoxDetect, except that their numbers of kernels per layer are 24, 48, and 48 respectively.

The first fully connected layer contains 64 hidden units, followed by a ReLU. Lastly, the second fully connected layer maps those 64 hidden units to 15 output units followed by a softmax nonlinearity corresponding to 14 species (shown in Figure 5) and an “other” (i.e. out-of-vocabulary) species class. The second fully connected layer also maps its 64 hidden units to 5 output units followed by a softmax nonlinearity corresponding to 4 families (shown in Figure 5) and an “other” family class, and single output unit followed by a sigmoid nonlinearity corresponding to Passeriformes or non-Passeriformes (order-level classification).

We note that there are no guarantees that the outputs of the model are hierarchically consistent; for example, the classifier can simultaneously predict *Cardinalidae* at the family level and *White-throated sparrow* at the species level even though white-throated sparrows are not cardinals. Since we do not have any guarantee of hierarchically consistency, we propose a method for selecting candidates which have this guarantee. Hierarchical consistency could be incorporated directly in the model by modeling joint class likelihoods instead of marginal class likelihoods, but we leave this question as future work.

### Hierarchical consistency

A simple method for selecting class candidates from the classifier output probabilities is to select the class with the largest output probability for each level in our taxonomy; however, this does not ensure that these class candidates are hierarchically consistent. In order to improve the robustness of the multilevel taxonomic classifier, we propose a method to ensure (top-down) *hierarchical consistency* for predictions. We define a procedure that, from a set of output probabilities for each taxon, produces class candidates that are hierarchically consistent. First, we select the class for the coarsest taxon (order, in this case) that has the largest output probability. If this probability is greater than a threshold of 0.5, we select this class as the taxon’s candidate and select “other” otherwise. Then, for each subsequent taxon, we select the class with the largest output probability that is also a taxonomic child of the previous taxon’s candidate. If this probability is greater than a threshold of 0.5, then we select this class for this taxon’s candidate and select “other” otherwise. Once we obtain a candidate for the finest taxon, we complete the collection of class candidates for each taxon.

### Data curation

To train and validate BirdVoxClassify, we present an updated version of the BirdVox American Northwest Avian Flight Call Classification (BirdVox-ANAFCC, or ANAFCC for short) Dataset [52], which we refer to as ANAFCC-v2. This dataset aggregates isolated flight calls from different data sources: BirdVox-full-night, CLO-43SD, CLO-SWTH, CLO-WTSP [**?**], the Macaulay Library [**?**], Xeno-Canto [**?**], and Old Bird [**?**]^6^. An expert ornithologist (AF of the authors) verified and re-annotated each clip and aligned each flight call precisely at the center of its corresponding clip. We map the resulting annotations onto our taxonomy as shown in Figure 5. This new version of ANAFCC, v2.0, contains additional flight calls from full-night which did not appear in the initial release, v1.0.

In order to better match our heterogenous development set to data found in realistic acoustic monitoring scenarios, we create training and validation subsets by finding a suitable partition of the ANAFCC data sources that is appropriately sized and have species distributions similar to that of the 296h dataset. To do this, we first pose the task of allocating data sources to the validation set as a knapsack problem [54] where we treat individual data sources as items. In the case of full-night we also treat clips from different recording units as separate sources. Each item has a weight corresponding to the number of annotated audio clips from the data source contains. We set the knapsack size according to our desired validation set size and the find the optimal knapsack using the dynamic programming algorithm implemented in Google OR-Tools [55]. We obtain optimal knapsacks for knapsack sizes corresponding to between 15–30% of the total number of examples, giving us a candidate set of appropriately sized validation subsets.

Given the data sources of a validation subset, we map all to the corresponding training subset. Finally, from this set of appropriately sized candidate partitions, we select the partition where the species distribution of both subsets are most similar to that of 296h. More precisely, we choose the partition with the lowest average Jensen-Shannon divergence between the species distributions of the split subsets and 296h.

### Training

We train the model to minimize a uniformly weighted summation of categorical cross-entropy for the species-level outputs, categorical cross-entropy for the family-level outputs, and binary cross-entropy loss for the order-level output. This multitask training method presented in prior species classification research [52] improves species classification performance over species-only training. We train the models using the Adam optimizer with initial learning rate set to 10^−4^. We also apply *L*^2^ regularization on the synaptic weights of the linear layers, using a multiplicative factor of 10^−5^ for the first linear layer and using a factor of 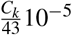 for each output layer for level *k* of the taxonomy with *C*_*k*_ classes. The output layer regularization factor is chosen so that each synaptic weight for the output layer is the same as in the original method [36].

### Evaluation

To evaluate BirdVoxClassify, we present a new version of the BirdVox 14 Species Dataset (BirdVox-14SD) [52], which we refer to as BirdVox-14SD-v1.1. A derivative of BirdVox-300h, BirdVox-14SD comprises roughly 14k isolated clips of flight calls alongside their human annotations, which are mapped to taxonomy shown in Figure 5. All clips last 500 ms and are centered around the annotated flight call. In comparison with the previous version (v1.0), the updated version (v1.1) addresses some edge cases regarding the alignment of clip boundaries.

We evaluate the predictions with and without enforcing hierarchical consistency to understand the impact of our hierarchical consistency procedure. We use F1 score for each class and overall to evaluate classifier performance, as shown in Figure 11.

### Results

With our hierarchical consistency procedure, the classifier achieves an F1 score of 72.82% whereas it achieves an F1 score of 66.71% without our hierarchical consistency procedure. We also see that for each class, enforcing hierarchical consistency results in matched or improved performance.

Figure shows the 10 confusion matrix between predicted classes and the ground truth in BirdVox-14SD-1.1. We observe that this matrix has a block structure: most of the off-diagonal confusions level correspond to different species of the same taxonomical family.

### Example use case: Swainson’s Thrush *(Catharus ustulatus)*

The sections above have focused on the evaluation of individual components in BirdVox: sensor fault detection, flight call detection, and species classification. It remains to be seen how BirdVox operates once these elements are integrated within a given application. For this purpose, we run the complete BirdVox pipeline on all recordings from the BirdVox-full-season dataset. This dataset contains 6,671 hours of audio; which, given a hop length of *τ* =15 ms, translates to

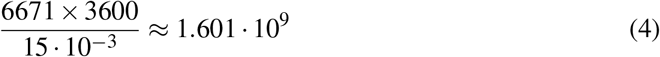

instances of Fast Fourier Transform (FFT). The convolutional neural network in BirdVoxDetect predicts event detection function at a rate of 20 Hz, hence a total of 6671 × 3600 × 20 ≈ 4.803.10^8^ > predictions. Furthermore, the number of synapses in the first layer of BirdVoxDetect is equal to 128 × 104 × 24 ≈ 2.995 × 10^5^. Because each synapse is encoded over 32 bits, or four bytes, the throughput of our computation is at least

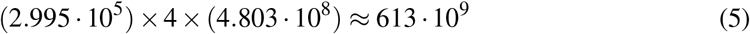

bytes, or 613 terabytes. Lastly, the output contains 6671 × 3600 × 20 × 4 ≈ 1.921 10^9^ bytes; i.e., around two gigabytes. These numbers demonstrate the need for parallel computing in the analysis of the full-season dataset. For this reason, we use the high-performance computing facility of New York University^7^ and parallelize execution massively over hundreds of CPU cores. In this way, the computation completes within a few hours.

Figure 13 illustrates the flight call activity for one of the most vocal species in the dataset: Swainson’s Thrush *(Catharus ustulatus)*. We express this flight call activity as the average number of flights calls across all non-faulty sensors in the network on a given day. Because sensor faults appear intermittently during the migration season (see 6), this average is not necessarily taken over the same subset of sensors between one day and the next. With that caveat in mind, we observe some consistent trends: busy migration activity on Sep. 13^th^ and 14^th^, followed by a decrease to almost zero detected flight calls, followed by a large peak on Sep. 23^th^. Juxtaposing these trends with a chart of crowdsourced observations on the ground (Figure 13, bottom) confirms that these trends are ecologically meaningful for the months of September.

**Figure 13.**
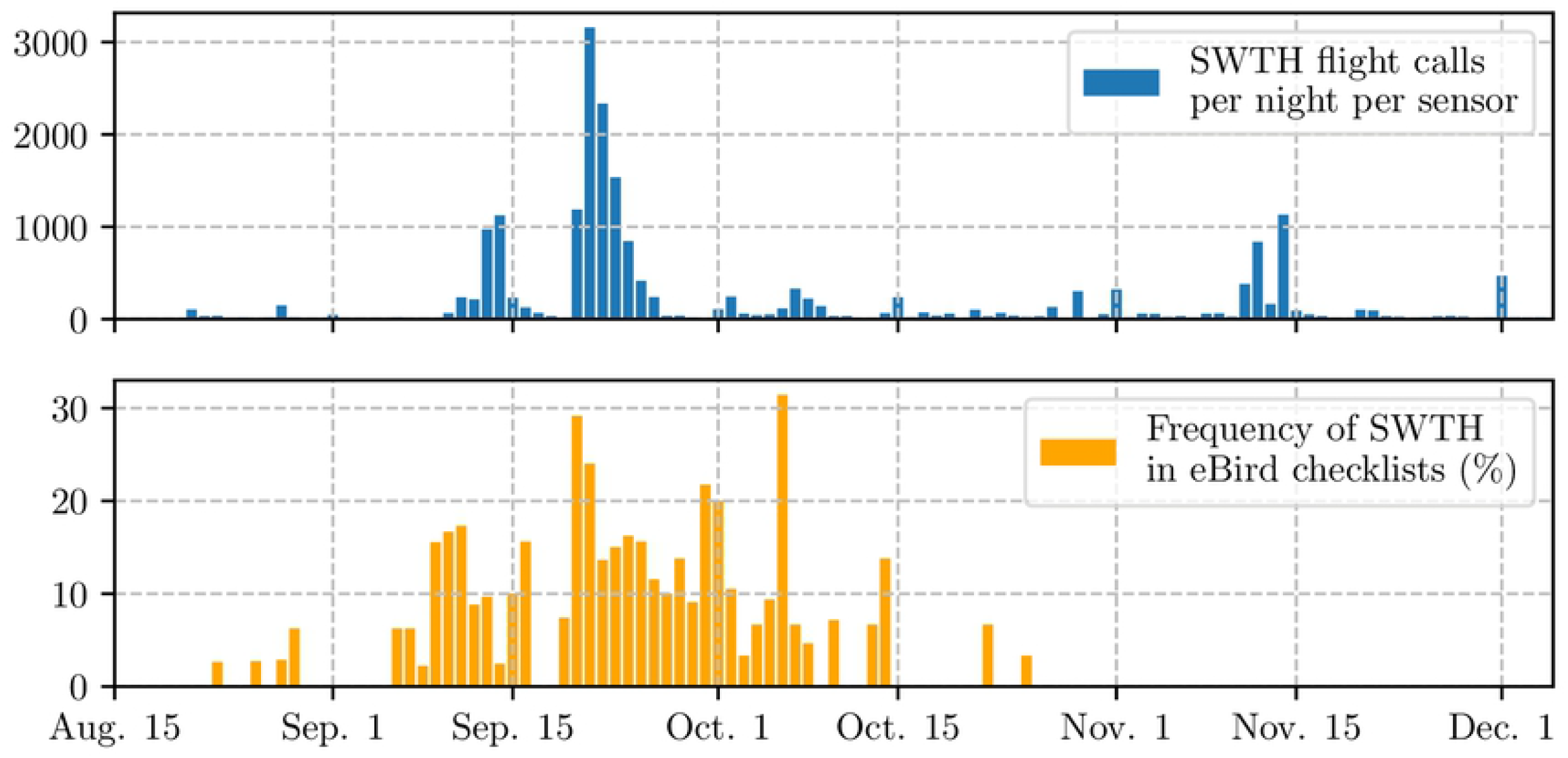
Comparison between machine listening and crowdsourced observations. Top: density of nocturnal flight calls from the Swainson’s Thrush *(Catharus ustulatus)* between midnight and 6 a.m. during the fall 2015 migration season in Tompkins County, NY, USA. Direct estimation from the BirdVox machine listening system on the full-season dataset (6,671 hours from nine sensors). Bottom: frequency of occurence of the Swainson’s Thrush among eBird checklists in Tompkins County for the same period.

However, we also notice that BirdVox predicts a relatively large flight call activity of Swainson’s Thrush in November and even December. Yet, domain-specific knowledge about the migration cycle of this species informs us that such a prediction is spurious, and is the result of mislabeling by the machine listening system. This observation indicates that BirdVox, despite being the current state of the art in flight call detection and classification, does not suffice on its own to monitor bird migration. Instead, it must be paired with radar measurements and/or crowdsourced observations so as to bring value for data-driven research in population ecology and conservation science. We leave this question to future work.

## Conclusion

The emerging field of machine listening for bird migration monitoring has the potential to elucidate some long-lasting questions in avian population ecology and inform conservation science efforts. In this paper, we have presented BirdVox, a cyber-physical system which is capable of detecting and classifying over fourteen species of flight calls from an acoustic sensor network on the ground. BirdVox integrates state-of-the-art components in signal processing and machine learning, such as per-channel energy normalization (PCEN) and deep convolutional neural networks (CNN). It also features novel elements such as a sensor fault detector trained with active learning and a rule-based algorithm for “hierarchical consistency” in the classification of living organisms. Our paper has shown that, once all elements are composed, BirdVox produces a daily log of flight call counts that, in the case of the most vocal species (Swainson’s Thrush), aligns with observations on the ground. We have released the automatic detector of BirdVox as part of an open-source software library, named BirdVoxDetect. Since this release, a community of flight call enthusiasts has adopted the tool and is currently using it to ease the process of nocturnal bird migration monitoring.

Beyond the technical aspects of BirdVox, it is worth stressing that the problem of flight call monitoring encompasses eleven orders of magnitude in terms of time scales: from a few microseconds for a digital audio sample up to millions of seconds for a full season. Figure 14 illustrates some of these time scales. Meanwhile, prior research on machine listening for the detection and classification of flight calls was carried out over four or five orders of magnitude: that is, up to one to ten seconds of time scale at most. With this paper, we aim to fill this gap in research by providing large-scale open audio datasets with expert annotation: BirdVox-full-season, BirdVox-296h, and BirdVox-14SD-v1.1. In future work, we will evaluate whether pairing machine listening with radar measurements and/or crowdsourced observations benefits the species-specific monitoring of bird migration at the macroscopic scale.

**Figure 14.**
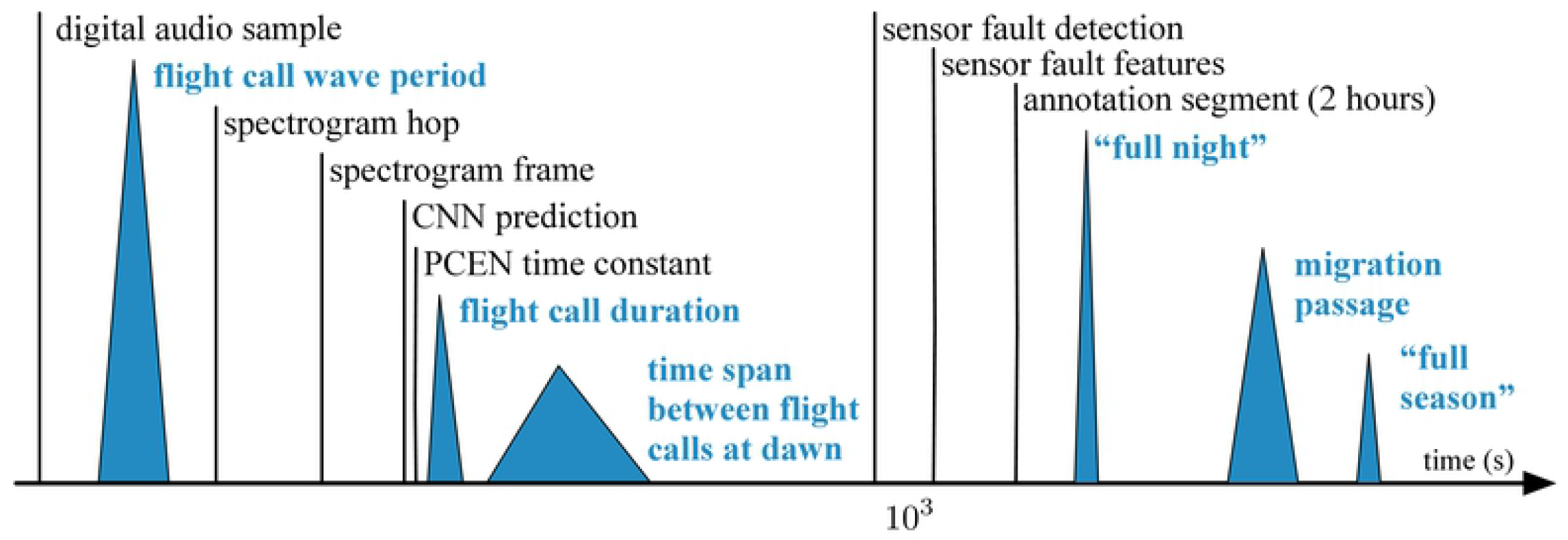
Timescales of bird migration monitoring with bioacoustic sensor networks. Blue triangles represent natural time scales, whereas black vertical lines represent our design choices.

## Acknowledgments

This work is supported by NSF awards 1633259 and 1633206, the Leon Levy Foundation, a Google faculty award, and an Atlanstic2020 project on “Trainable Acoustic Sensors” (TrAcS). We thank Jessie Barry, Ian Davies, Tom Fredericks, Jeff Gerbracht, Sara Keen, Holger Klinck, Anne Klingensmith, Ray Mack, Peter Marchetto, Ed Moore, Matt Robbins, Ken Rosenberg, and Chris Tessaglia-Hymes for designing autonomous recording units and collecting data. We acknowledge that the land on which the data were collected is the unceded territory of the Cayuga nation, which is part of the Haudenosaunee (Iroquois) confederacy.

## Footnotes

1 Data repository of BirdVox-full-season: https://zenodo.org/record/5791744

2 For more information on the design of bioacoustic sensors for bird migration monitoring, visit: http://www.oldbird.org

3 Official website of Raven: https://ravensoundsoftware.com.

4 Data repository of BirdVox-296h: https://zenodo.org/record/4603643

5 Documentation of pescador: https://pescador.readthedocs.io

6 Official website of Old Bird, Inc.: https://www.oldbird.org

7 Link: https://sites.google.com/nyu.edu/nyu-hpc

